# Theoretical Expression for the Evolution of Genomic Island of Speciation: From Its Birth to Preservation

**DOI:** 10.1101/545103

**Authors:** T. Sakamoto, H. Innan

## Abstract

Ecological speciation could be driven by divergent selection that works to maintain phenotypes that are adaptive to each niche. In its early stages, genetic divergence (or F_ST_) can be maintained around the target sites of divergent selection, while in other regions, genetic variation can be mixed by gene flow or migration. Such regions of elevated genetic divergence are called genomic islands of speciation. In this work, we theoretically consider the evolutionary process of a genomic island of speciation, from its birth to stable preservation. Under a simple two-population model, we use a diffusion approach to obtain analytical expressions for the probability of initial establishment of a locally adaptive allele, the reduction of genetic variation due to the spread of the adaptive allele, and the process to the development of a sharp peak of divergence. Our result would be useful to understand how genomes evolve through ecological speciation with gene flow.

---

Agenomic island of speciation arises in the earlier stages of ecological speciation with gene flow (Wu 2001; Turner *et al.* 2005; Nosil 2012). Speciation can be initiated by the initial establishment of a locally adapted allele to a certain subpopulation. This local establishment could be stably maintained by divergent selection when the allele confers sufficient benefit in the adaptive subpopulation(s), but not (or even deleterious) in others. Due to recombination, the genomic regions that are affected by divergent selection is limited, thereby creating a peak of divergence along chromosome, that is, a genomic island of speciation. We are here interested in the evolutionary behavior of a genomic island of speciation, from its initial establishment to stable preservation.

Theoretically, it would be convenient to consider the process by dividing into three phases, the establishment, erosion and equilibrium phases, as illustrated in Figure 1. We consider a simple situation with two subpopulations, I and II. Assuming a relatively high migration rate between them, the levels of polymorphism within the two subpopulations are similar to each other (measured by the average numbers of pairwise nucleotide differences, *π*_*w*1_ and *π*_*w*2_, for subpopulations I and II). In the meantime, the population divergence (measured by *π*_*b*_, the average numbers of pairwise nucleotide differences between the two subpopulations) is very low (Figure 1A). Then, a *de novo* mutation (the star in Figure 1A) arises in subpopulation I, in which the mutation is advantageous but maladaptive (or deleterious) in subpopulation II. In the establishment phase, the mutation spreads in subpopulation I and nearly fixes (Figure 1B), but its frequency in subpopulation II is low because it should be selected against if migrated into subpopulation II. In a strict sense, this is not a fixation that can be mathematically treated as an absorbing state, because migration keeps providing maladaptive alleles. Therefore, after Kimura (1954), we hereafter use the terminology of “quasi-fixation” for this nearly fixed state. The quasi-fixation should occur quickly, and a selective sweep occurs in subpopulation I (Figure 1B), thereby creating an initial island. The initial genomic island should be as large as the region where genetic variation within the genomic island should be very low in subpopulation I, whereas genetic variation in subpopulation II may not be much affected by the selective sweep (Figure 1B). The erosion phase starts after the initial establishment, during which the genomic island gradually shrinks over time by recombination and migration (Figure 1C). Then, at the end, the genomic island appears as a stable sharp peak of divergence in the equilibrium phase (Figure 1D). The equilibrium size of the genomic island is mainly determined by the balance between selection intensity and the rates of recombination and migration.

**Figure 1.**
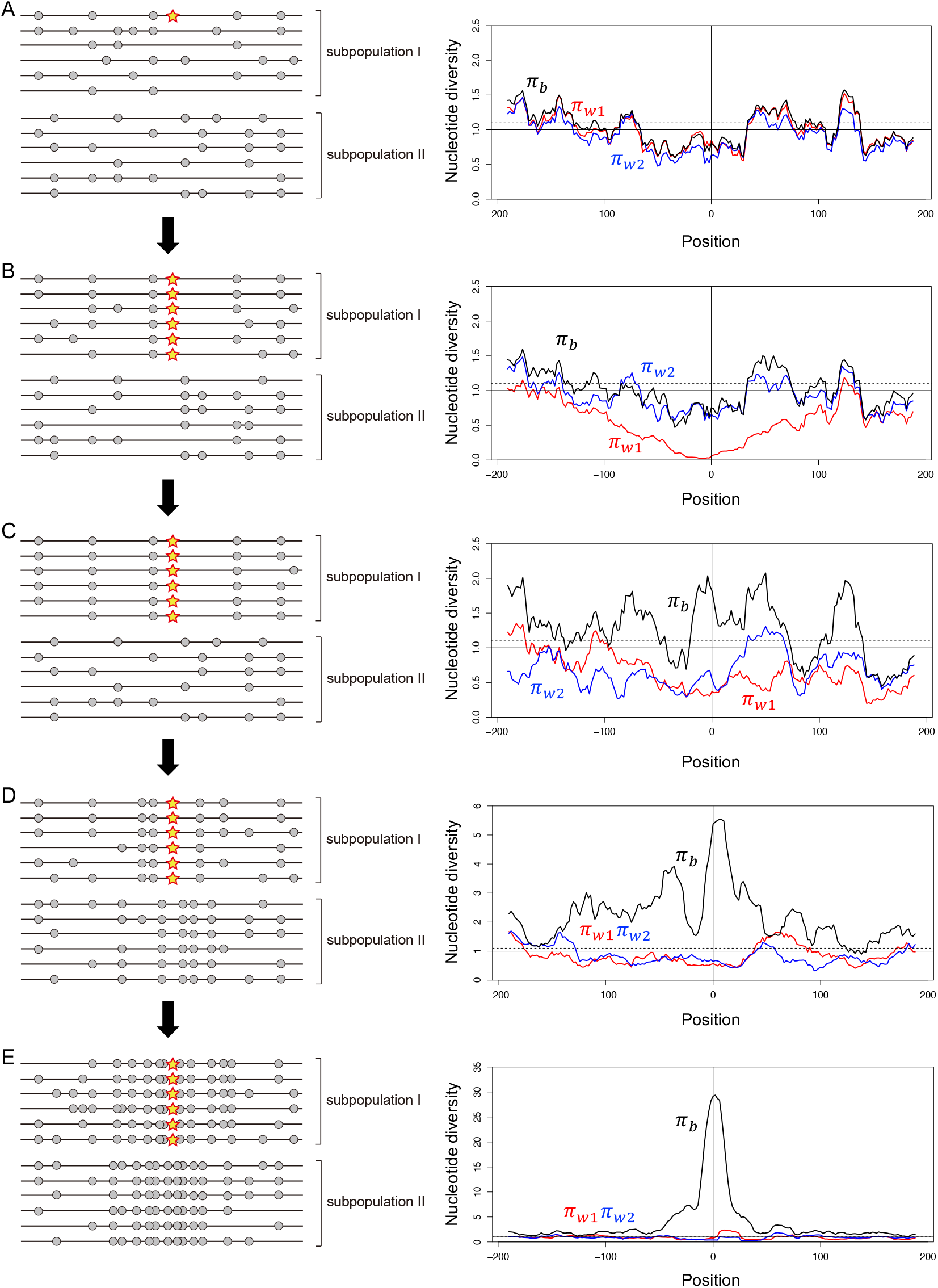
Illustrating the evolution of a genomic island of speciation in a simple two-population model with fairly high migration between them. (A) A locally adaptive de novo mutation arises in subpopulation I at position 0. A typical pattern of polymorphism is shown in left. The star is the locally adaptive mutation and gray circles are neutral polymorphism in the surrounding region. The right panel shows the spacial distributions of nucleotide diversity obtained by a simulation. The polymorphism levels within the two populations (*π*_*w*1_ and *π*_*w*2_) are in red and blue, and divergence between the two populations (*π*_*b*_) is in black. The y-axis is adjusted such that *E*(*π*_*w*1_) = *E*(*π*_*w*2_) = 1 under neutrality (the solid line), and the broken line exhibits *E*(*π*_*b*_). The entire simulated region is 400 kb if the population recombination rate = 0.001 per site is assumed. (B) The mutation quasi-fixes in subpopulation I, causing a drastic reduction in *π*_*w*1_. (C) Migration shuffles polymorphisms in the two subpopulations, while selection works to maintain the quasi-fixation of the mutation. (D) The divergence gradually increases around the mutation, and (E) a clear peak of divergence arises.

The scope of this work is to provide a unified and comprehensive theoretical understanding of the evolution of a new genomic island of speciation, from its birth to stable preservation in equilibrium. We use a simple two-population model, where migration is allowed between subpopulations I and II. Suppose a de novo mutation arises that confers a selective advantage specific to subpopulation I, which is the initial state of our system. Under this model, we derived the following:

for the establishment phase,

i. The establishment probability of the de novo mutation, that is, the probability that the mutation quasi-fixes in subpopulation I.
ii. The expected reduction of genetic variation within subpopulations I and II after the quasi-fixation (i.e., local sweep). for the erosion phase,
iii. The expected erosion of the initial island as a function of time since the quasi-fixation. and for the equilibrium phase,
iv. The expected shape of the genomic island at equilibrium.

There have been several theoretical works that focused on a specific part of these aspects. For (i) the established probability, perhaps the most flexible, useful theoretical framework was introduced by Barton (1987) in a general multiple-island-model. By using a diffusion approximation, Barton (1987) derived a partial differential equation for the establishment probability. Essentially the same result was obtained by Pollak (1966), who used a branching process and the establishment probability was derived from the probability generating function. Barton’s differential equation was solved and closed forms of the establishment probability have been available only in several specific situations in *continuous* habitat models. In a one-dimensional continuous habitat model, Barton (1987) solved his partial differential equation analytically assuming two forms of fitness gradient (linear and pocket). Kirkpatrick and Peischl (2013) used a branching process, from which they obtained a partial differential equation that is similar to that of Barton (1987). Then, the authors successfully incorporated changes in fitness gradient along time, and derived an approximate establishment probability.

In *discrete* population models, Barton’s general formula (and also Pollak’s one) is difficult to handle and have not been fully explored even in a simple two-population model with symmetric migration. Therefore, the currently available theoretical results are not based on Barton’s differential equation, and have some limitations. In a continent-island model with *unidirectional* migration, Aeschbacher and Bürger (2014) solved the establishment probability of a locally beneficial mutation linked to another locally beneficial mutation that was already established, where mathematical treatment is quite straightforward because of unidirectional migration (see also Yeaman *et al.* 2016). Yeaman and Otto (2011) obtained an approximate establishment probability by using a heuristic approach that is a combination of the leading eigenvalue of the transition matrix of deterministic process and Kimura’s formula of fixation probability (Kimura 1962). As shown in their paper, this formula well describes the establishment probability when a de novo mutation arises in the adapted subpopulation (i.e., subpopulation I in our model), but it does not work when it arises in the maladapted subpopulation (i.e., subpopulation II in our model). Recently, Tomasini and Peischl (2018) provided an approximate established probability by assuming a slightly supercritical branching process. Their formula works well under the assumption of slightly supercritical approximation, namely, the leading eigenvalue of the transition matrix of deterministic model is not large, but it may not work well when the selection intensity in the adapted subpopulation is very large.

In this work, we derive a closed form formula of the establishment probability in a two-population model with bidirectonal migration along the formulation of Barton (1987). We extended Barton’s derivation with simultaneous quadratic equations and solved them allowing unequal subpopulation sizes. Our formula is more general than previous ones (Yeaman and Otto 2011; Tomasini and Peischl 2018); it works with strong selection and it allows that a de novo mutation can arise either subpopulation I or II.

To our best knowledge, there is no theoretical work on the hitch-hiking process of a local sweep in a two-population model. With regard to a single population model, many studies theoretically investigated the reduction of polymorphism due to a selective sweep. These studies considered a selected site and a linked neutral site, and assumed that a very advantageous mutation arises and goes to fixation in the population. Along this fixation, they derived how much polymorphism can be reduced at the linked site. Maynard Smith and Haigh (1974) first obtained the reduction of polymorphism, where the stochastic effect of genetic drift at the linked site was ignored. The model was extended to include the stochastic effect by using a coalescent approach (Kaplan *et al.* 1989) and by using a diffusion method (Stephan *et al.* 1992). It is worthy to note that Stephan *et al.* (1992) derived a nice analytical approximate formula (see also Barton 1998; Etheridge *et al.* 2006). Durrett and Schweinsberg (2004) used a different approach for a faster approximate simulation of a selective sweep and derived some analytical expressions (see also Schweinsberg and Durrett 2005).

There are several theoretical studies on a sweep in multi-population models available, but these considered a fixation across multiple subpopulations, not a local fixation. In a model with multiple subpopulations, Slatkin and Wiehe (1998) considered the process where a beneficial mutation fixes in the entire population through weak migration. Santiago and Caballero (2005) considered a two-population model with a more general initial state and derived analytical expressions under the assumptions of weak migration. Kim and Maruki (2011) allowed stronger migration and derived analytical expression in a two-population model. Our interest is different from these studies in that we consider a locally beneficial mutation that can quasi-fix only in the adaptive subpopulation (not the entire population). We here extended the theory of Stephan’s diffusion model (Stephan *et al.* 1992) to a two-population model, and considered how much polymorphism can be reduced at a linked site after a local sweep.

The common interest in the erosion phase is how a genomic island decays along time. We here consider this process after a local sweep as described in Figure 1. A local sweep creates a “block” of a fairly long region with almost no genetic variation in the adaptive population (i.e., subpopulation I in our model). In this work, given an arbitral configuration of genetic variation after a local sweep, we analytically obtain the moments of allele frequency at a linked site, with which we describe how a genomic island decays. Yeaman *et al.* (2016) investigated a similar problem in a different situation, where a secondary contact occurred between already diverged populations. In their model, erosion starts when there already are a large number of fixed sites that spread over the genome, and islands appear because selection works to maintain divergence at selected site(s), while losing divergence in other regions through homogenization by migration. By using the structured coalescent, they obtained the expected spacial distribution of F_ST_ (in terms of relative coalescent time) around a selected site as a function of the time since the secondary contact. They also considered the scenario where a de novo mutation creates a genomic island, but their derivation did not consider the effect of selective sweep of the de novo mutation, which may be slightly unrealistic. It should be noted that, because our derivation accepts any arbitral initial allele frequency at a linked site, it can be applied to any situation, not only that after a secondary contact but also that after a local sweep.

In the equilibrium phase, the balance between selection, migration, recombination and mutation holds. Theoretical treatment at equilibrium is relatively straightforward, and there are several theoretical studies on the spacial distribution of F_ST_ (Charlesworth *et al.* 1997; Akerman and Bürger 2014; Yeaman *et al.* 2016). Under our framework for the erosion phase, essentially the same result can be provided as a special case with time going to infinity.

## MODEL

We consider a random mating two-population model with discrete generation, which follows the Wright-Fisher reproduction. The diploid population sizes of subpopulations I and II are assumed to be constant at *N*_1_ and *N*_2_, respectively. As illustrated in Figure 1, we are specifically interested in selection for local adaptation in subpopulation I. We consider a genomic region encompassing a selected site at position 0, which is referred to as locus A (Figure 2). At locus A, two alleles (A/a) are allowed with no recurrent mutation between them. Allele A confers a selection coefficient *s*_1_ in subpopulation I and *s*_2_ in subpopulation II (we assume *s*_1_ > 0 and *s*_2_ < 0). Additive selection is assumed so that the fitness of individuals with AA, Aa and aa are given by 1 + 2*s*_1_, 1 + *s*_1_ and 1 in subpopulation I, and 1 + 2*s*_2_, 1 + *s*_2_ and 1 in subpopulation II. Selection works only at this selected site, and all remaining sites are assumed to be neutral. For the following derivation under a two-locus model, we consider a secondary neutral site (locus B), at which two alleles (B/b) are allowed with recurrent mutation between them (Figure 2). The mutation rate from B to b is *u* and that from b to B is *v*. *r* is the recombination rate between the two loci, A and B.

**Figure 2.**
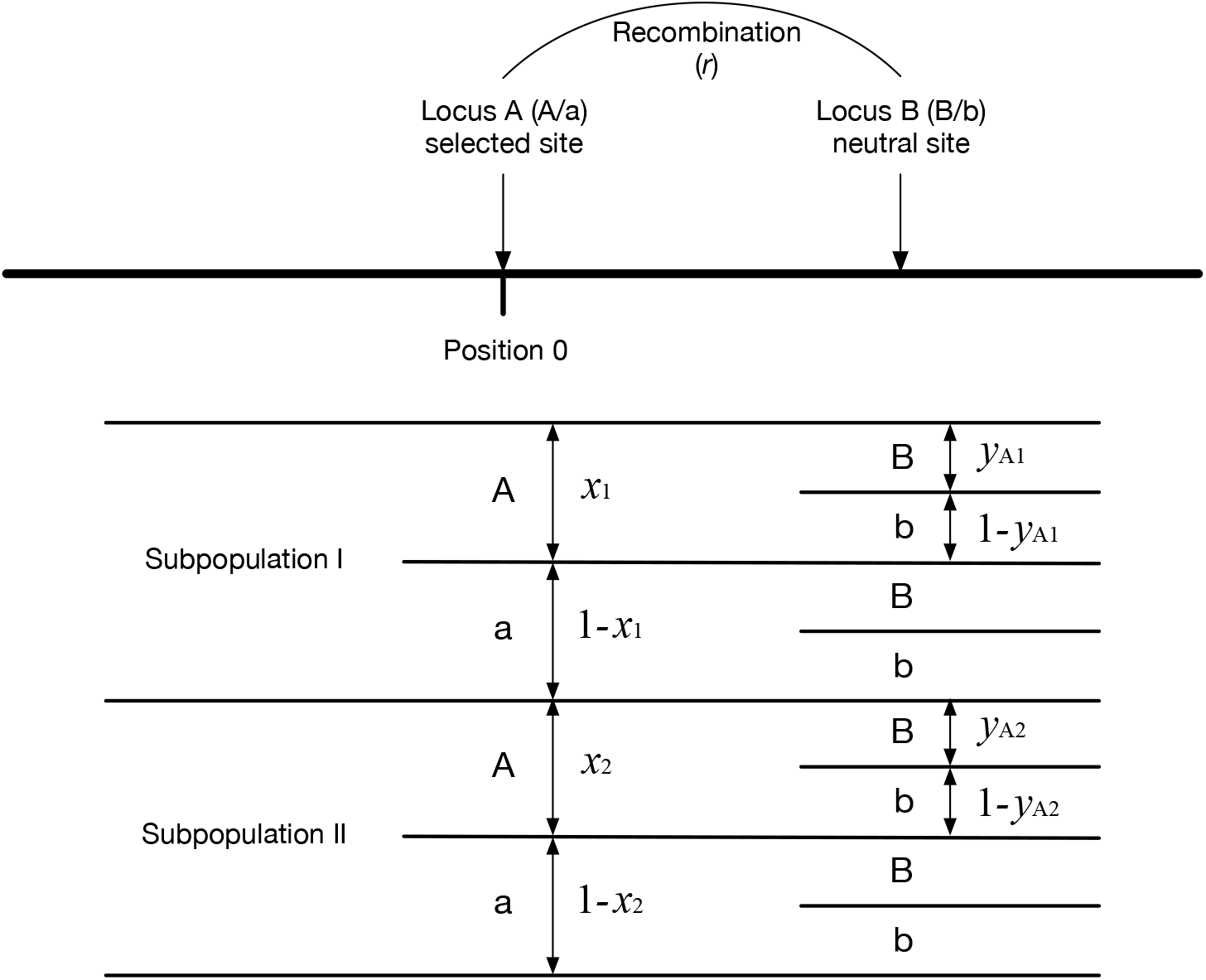
Two-locus model used in this work.

The system starts when a de novo mutation (allele A) arises in a single individual either in population I or II, where allele a is fixed in both subpopulations. Therefore, the initial state is (*x*_1_, *x*_2_) = (1/2*N*_1_, 0) or (0, 1/2*N*_2_), where *x*_1_ and *x*_2_ are frequencies of the new allele A in subpopulations I and II, respectively. Throughout this article, we assume strong selection and weak migration so that maladapted individuals are rare in each subpopulation once the initial establishment is achieved.

### Establishment probability

We derive the establishment probability of a new de novo allele using the general framework of Barton (1987), who derived a simultaneous quadratic equation from the diffusion theory. This section focuses only on the selected locus A (see Figure 2), at which we are interested in the probability that allele A quasi-fixes in subpopulation I. Following previous studies (Haldane 1927), we consider that the establishment probability can be essentially obtained as the probability that the new mutation increases in frequency and escapes from immediate extinction. This is because, with the assumption of strong selection, the behavior of such an allele is almost deterministic once it escapes from extinction by genetic drift.

Let *u*(*x*_1_, *x*_2_) be the establishment probability when the frequencies of allele A are *x*_1_ and *x*_2_ in the two subpopulations. By using an analogous procedure to Barton (1987), we derive *p*_1_ = *u*(1/2*N*_1_, 0) and *p*_2_ = *u*(0, 1/2*N*_2_), the establishment probability when the new allele arises in subpopulations I and II, respectively. According to the diffusion theory, *u* satisfies the Kolmogorov backward equation:

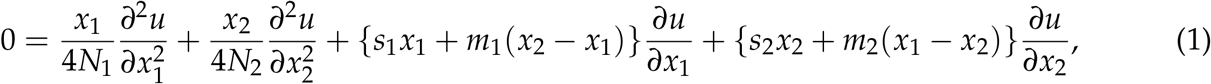

where *m*_1_(*m*_2_) is the proportion of immigrant individuals just after migration in subpopulation I (II). To keep the subpopulation sizes constant, we assume *N*_1_*m*_1_ = *N*_2_*m*_2_, and we ignore higher order terms of 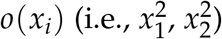. This is reasonable because of the assumption that the establishment probability is mainly determined at low frequencies. Because the extinction probability of each individual mutation is independent to each other, we can write *u* as

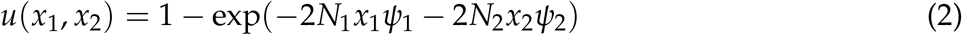

where exp(−*ψ*_*i*_) is the extinction probability of a new mutant in subpopulation *i*, therefore, *p*_*i*_ = 1 – exp(−*ψ*_*i*_). By substituting Equation 2 into Equation 1, we have

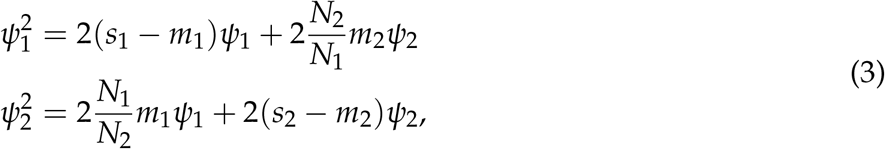

which corresponds to Equation 4b in Barton (1987). Then, the above equations deduce

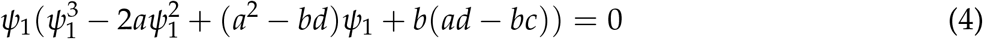

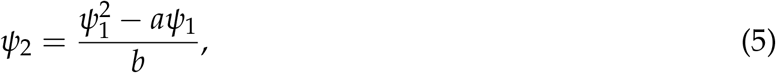

where 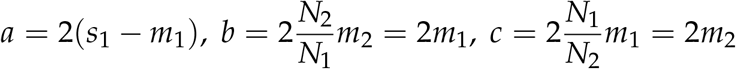, and *d* = 2(*s*_2_ – *m*_2_). Equation 4 can be solved by using the solution of cubic equation. Equations 4 and 5 have at most one solution which fulfills *p*_1_ > 0 and *p*_2_ > 0. The condition where Equations 4 and 5 have such a solution is *a* + *d* > 0 or *ad* – *bc* < 0, which corresponds to the situation where the deterministic growth rate of the mutant allele is positive (see APPENDIX A for details).

Figure 3 shows the establishment probability from Equations 4 and 5 as a function of migration rate. We first consider a symmetric model (*N*_1_ = *N*_2_ = 1000), and two selection intensities (*s*_1_ = 0.02 and *s*_1_ = 0.1) are assumed, while *s*_2_ = −0.01 is fixed (Figures 3A and B). The establishment probability can be computed when a locally adaptive mutation arises either in subpopulation I or II, represented as *u*(1/2*N*_1_, 0) and *u*(0, 1/2*N*_2_), respectively. We performed a forward simulation to check the performance of our analytical result. For each parameter set, we ran 1,000,000 independent replications of simulation and counted the number of replications where the new allele A was preserved in 10,000 generations. The establishment probability was then obtained as the proportion of such replications. Therefore, it includes replications where alleles A and a coexisted (case C) and those where A is completely fixed in both subpopulations (case F). The proportion of case C in the established replications (*Pc*) decreases with increasing the migration rate (see below).

**Figure 3.**
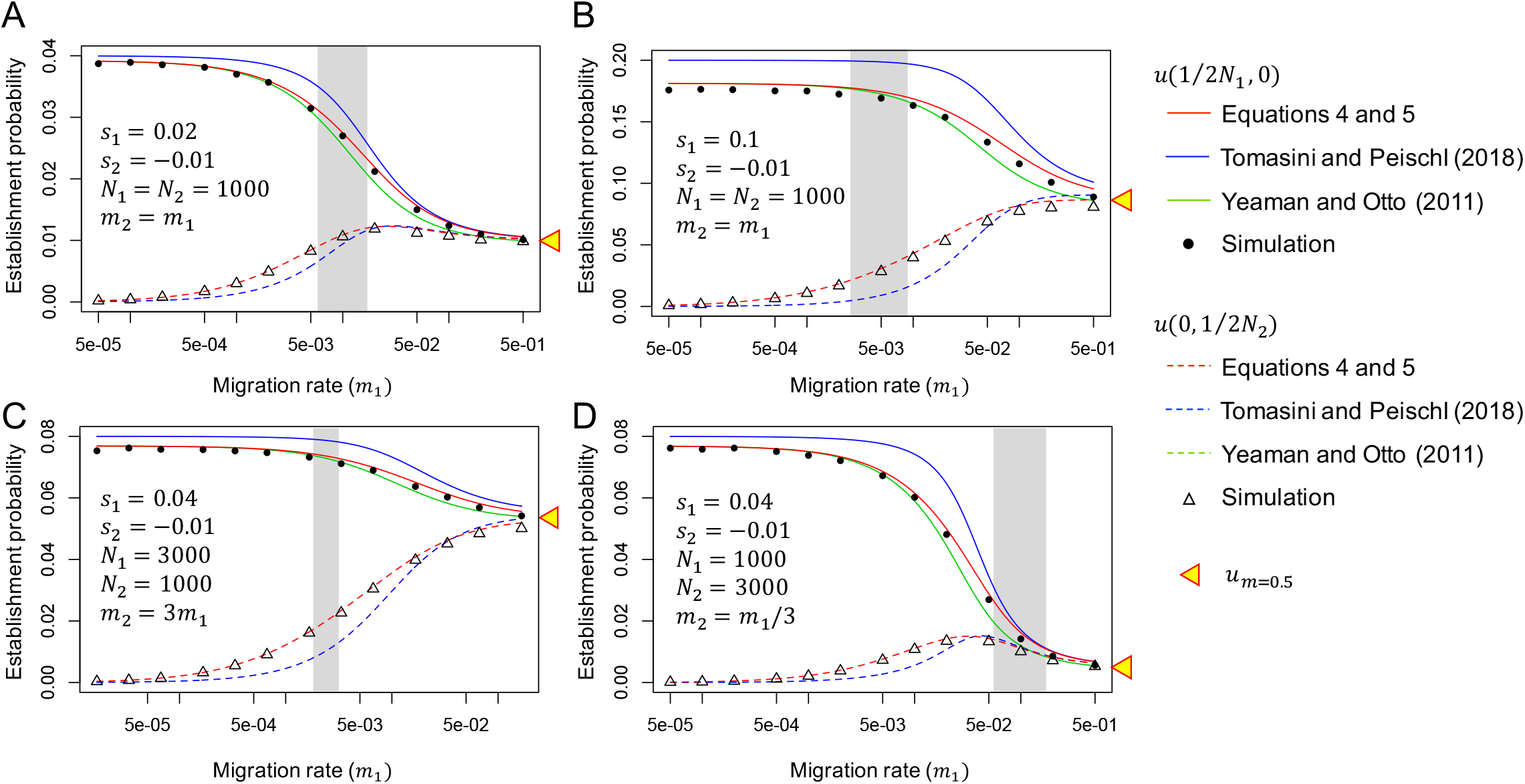
Establishment probability as a function of migration rate. (A) Weak selection (*s*_1_ = 0.02 and *s*_2_ = −0.01) and strong selection (*s*_1_ = 0.1 and *s*_2_ = −0.01) are assumed in a symmetric model (*N*_1_ = *N*_2_). (C, D) Asymmetric population settings are considered (*N*_1_ = 3*N*_2_ in C and *N*_1_ = *N*_2_/3 in D). Our result in red is compared with those of Tomasini and Peischl (2018) and Yeaman and Otto (2011), together with the result of our forward simulation. The establishment probability for a mutation that arises in subpopulation I (*u*(1/2*N*_1_, 0)) is shown by solid lines and closed circles, and that for a mutation that arises in subpopulation II (*u*(0, 1/2*N*_2_)) is shown by broken lines and open triangles. The establishment probability at the high migration limit (*m* = 0.5) is shown by a yellow triangle. In each panel in Figure 3, a gray region is placed such that *Pc* > 0.9 in the left, while *Pc* < 0.1 in the right.

Our result (red in Figure 3) is in an excellent agreement with the simulation result: *u*(1/2*N*_1_, 0) is approximately 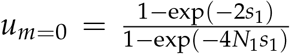 when the migration rate is very low, consistent with the prediction in a single population model (Kimura 1957). As the migration rate increases, *u*(1/2*N*_1_, 0) decreases and *u*(0, 1/2*N*_2_) increases, and they become similar to each other. With a very high migration rate (*m* ∼ 0.5), the two subpopulations can be considered as a single random-mating population, and the fixation probability of a single mutation is mainly determined by the average selection coefficient, 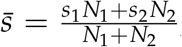, namely, 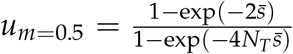 where *N*_*T*_ = *N*_1_ + *N*_2_ (Nagylaki 1980). Indeed, in our simulations, allele A was fixed in both populations in almost all established cases (*Pc* = 1). In each panel in Figure 3, a gray region is placed such that *Pc* > 0.9 in the left, while *Pc* < 0.1 in the right. It is indicated that the pattern dramatically changes in a short range of *m*_1_, and the left side is the scope of this article. Similar results were also obtained in asymmetric models (*N*_1_ = 3*N*_2_ in Figure 3C and *N*_1_ = *N*_2_/3 in Figure 3D).

Figure 3 quantitatively compares our analytical results with those of previous studies (Tomasini and Peischl 2018; Yeaman and Otto 2011). It is found that *u*(1/2*N*_1_, 0) from Yeaman and Otto (2011) is almost as good as ours, but unfortunately *u*(0, 1/2*N*_2_) was not provided by Yeaman and Otto (2011). It seems that Tomasini and Peischl (2018) overestimates *u*(1/2*N*_1_, 0) and underestimates *u*(0, 1/2*N*_2_).

### Reduction of genetic variation due to a selective sweep

When a new locally adaptive mutation (a→A) arises and quasi-fixes in subpopulation I, genetic variation in the surrounding region in subpopulation I should be dramatically reduced due to the hitch-hiking effect. In this section, we consider a two-locus model as defined in Figure 2. We derive the degree of reduction in heterozygosity at a linked neutral site (locus B) in subpopulation I, *D*_*local sweep*_, by extending the diffusion approach of Stephan *et al.* (1992), who investigated the effect of hitch-hiking in a single population model with no population structure.

### Overview of Stephan *et al.* (1992)

We first introduce the approach of Stephan *et al.* (1992) briefly, which provides the basis of our derivation below. The expected reduction of heterozygosity at locus B for a single population model with diploid size *N* is denoted by *D*_Stephan et al._. With the assumption of strong selection, Stephan *et al.* (1992) assumed that the behavior of the frequency (*x*) of the beneficial allele A with selection coefficient, *s*, follow a deterministic function:

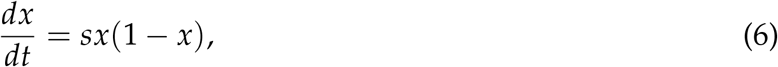

where selection is additive. It should be noted that *x* with no subscript denotes the frequency of A in the single population model, whereas in our two-population model, the frequencies of A in subpopulations I and II are denoted by *x*_1_ and *x*_2_, respectively (see Figure 2). We consider another biallelic neutral locus (B/b), and the recombination rate between between this neutral locus and the selected locus is assumed to be *r*. *y*_*A*_ is the frequency of B among A-chromosomes and *y*_*a*_ is the frequency of B among a-chromosomes. Then, the expected changes of an arbitrary function *f* (*y*_*A*_, *y*_*a*_) is described as the following ordinary differential equation:

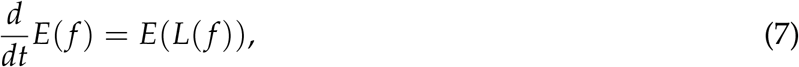

where *L* is a differential operator of the Kolmogorov backward equation:

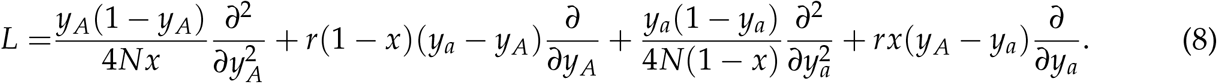

By using this formula, Stephan *et al.* (1992) solved the first and second moments of *y*_*A*_ and *y*_*a*_ after a sweep, from which the expected reduction of heterozygosity at the linked site can be computed numerically. With some approximation, Stephan *et al.* (1992) further obtained a nice closed form of the solution:

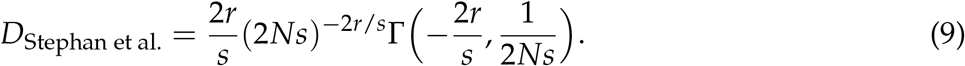

In this work, we found that this equation somehow undervalues the effect of random genetic drift perhaps due to the approximation of Stephan *et al.* (1992). It is known that heterozygosity decreases by genetic drift by a factor of 1/2*N* per generation. To correct for this factor, we obtain the expected reduction of heterozygosity along the fixation at the selected site as exp(−log(2*N*)/*Ns*), because the fixation time is approximately given by

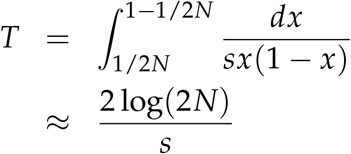

Then, we add this factor into Equation 9:

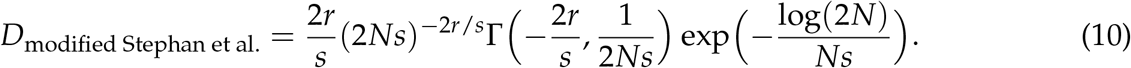

We found that this heuristic approach is in a very good agreement with the numerical solution obtained by directly computing Equation 8.

### Local sweep in the two-population model

In this work, we extend Stephan *et al.*’s derivation (1992) to the two-population model defined above (Figure 2). We first consider the dynamics of the new mutant allele frequency (*x*_1_) at the selected locus (position 0) in the subpopulation I. The major difference from the corresponding formula in Stephan *et al.* (1992) (i.e., Equation 6) is that the effect of migration should be considered in the two-population model. Because maladaptive allele A is very rare in subpopulation II under the assumption of strong selection and low migration, we can ignore migrants with A allele from subpopulations II to I. Then, the dynamics of *x*_1_ could be approximated by a deterministic function:

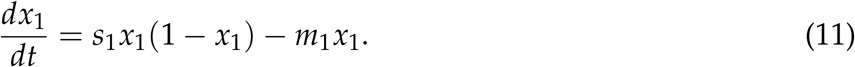

We set the time such that *t* = 0 when the mutation arises and *t* = *τ* when the mutation quasi-fixes. We next consider the neutral locus B (B/b). As illustrated in Figure 2, *y*_*A*1_ (*y*_*A*2_) is the frequency of haplotype A-B among A-chromosomes in subpopulation I (II), and *y*_*a*1_ (*y*_*a*2_) is the frequency of haplotype a-B among a-chromosomes in subpopulation I (II). We assume that *y*_*A*2_ is very small throughout the sweep process. Then, the expected changes of an arbitrary function *f* (*y*_*A*1_, *y*_*a*1_, *y*_*a*2_) is described as the following ordinary differential equation:

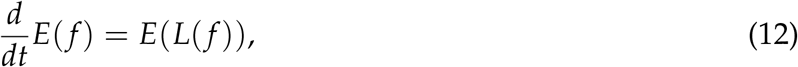

where *L* is a differential operator of the Kolmogorov backward equation. Following Ohta and Kimura (1969), we obtain *L* for our model as

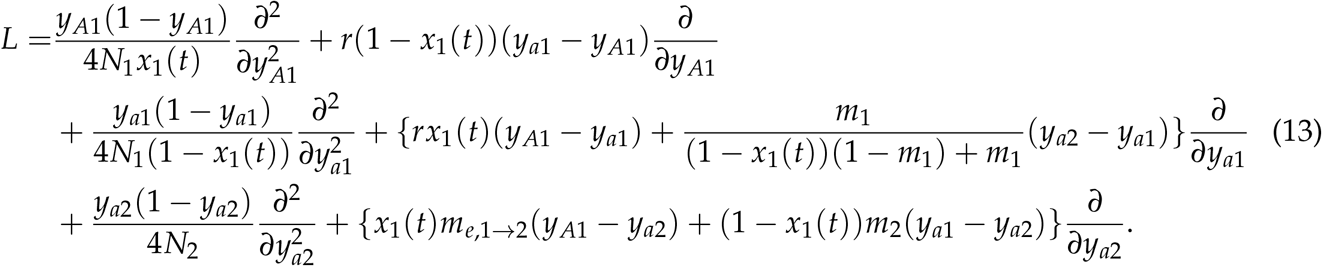

In this equation, the effect of selection on locus B is incorporated such that A-chromosomes will be selected out immediately if migrated from subpopulations I to II. In other words, with the linkage effect, the migration rate of A-chromosomes at the B locus is *effectively* reduced to *m*_*e*,1→2_:

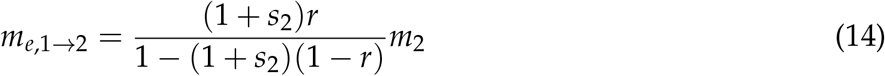

(Bengtsson 1985). Then, we can compute the first and second moments of *y*_*A*1_ and *y*_*a*2_ after the quasi-fixation of allele A (i.e., *y*_*A*1_(*τ*) and *y*_*a*2_(*τ*)), from which we can obtain heterozygosity within each subpopulations (*h*_*w*1_ and *h*_*w*2_) and between them (*h*_*b*_) at *t* = *τ* as

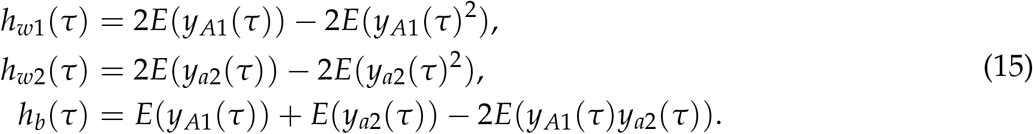

Then, the expected reduction of heterozygosity is obtained as

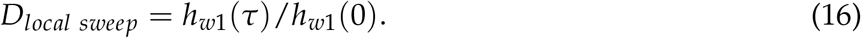

Generally, *D*_*local sweep*_ involves the initial frequencies, *y*_*a*1_(0) and *y*_*a*2_(0). However, it should be noted that their quantitative effect on *D*_*local sweep*_ is not large unless *y*_*a*1_(0) and *y*_*a*2_(0) are not very similar. Figure 4 shows the effect of migration on the reduction in heterozygosity. The plot in red is the case of no migration, where our result is essentially identical to Stephan *et al.* (1992). For the cases with migration, we assumed *N*_1_ = *N*_2_ = 1000, *m*_1_ = *m*_2_ = *m*. For each parameter set, filled circles represent the average over 100,000 replications of forward simulation. In Figure 4, *h*_*w*1_(*τ*), *h*_*w*2_(*τ*) and *h*_*b*_(*τ*) are plotted such that *h*_*w*1_(0) = *h*_*w*2_(0) = 1 before the sweep, so that *h*_*w*1_(*τ*) directly corresponds to *D*_*local sweep*_. In all cases, our theoretical result from Equation 13 is in excellent agreement with the simulation results. It is found that the effect of a sweep seems to be only on subpopulation I, and there is almost no effect on the variation in subpopulation II. As going further from the selected site at position 0, *D*_*local sweep*_ is larger for a higher migration rate. It is indicated that migration brings standing variation maintained in subpopulation II into subpopulation I, thereby increasing the polymorphism level in subpopulation I. We can observed a slight increase of *h*_*b*_(*τ*) around the selected site at position 0. If we assume 1 – *h*_*w*_(*τ*)/*h*_*all*_ (*τ*) roughly approximates F_ST_ where *h*_*all*_ is heterozygosity when the two subpopulations are merged together, it can be said that a local sweep creates a relatively wide region of elevated F_ST_, which can be considered as an initial genomic island of speciation.

**Figure 4.**
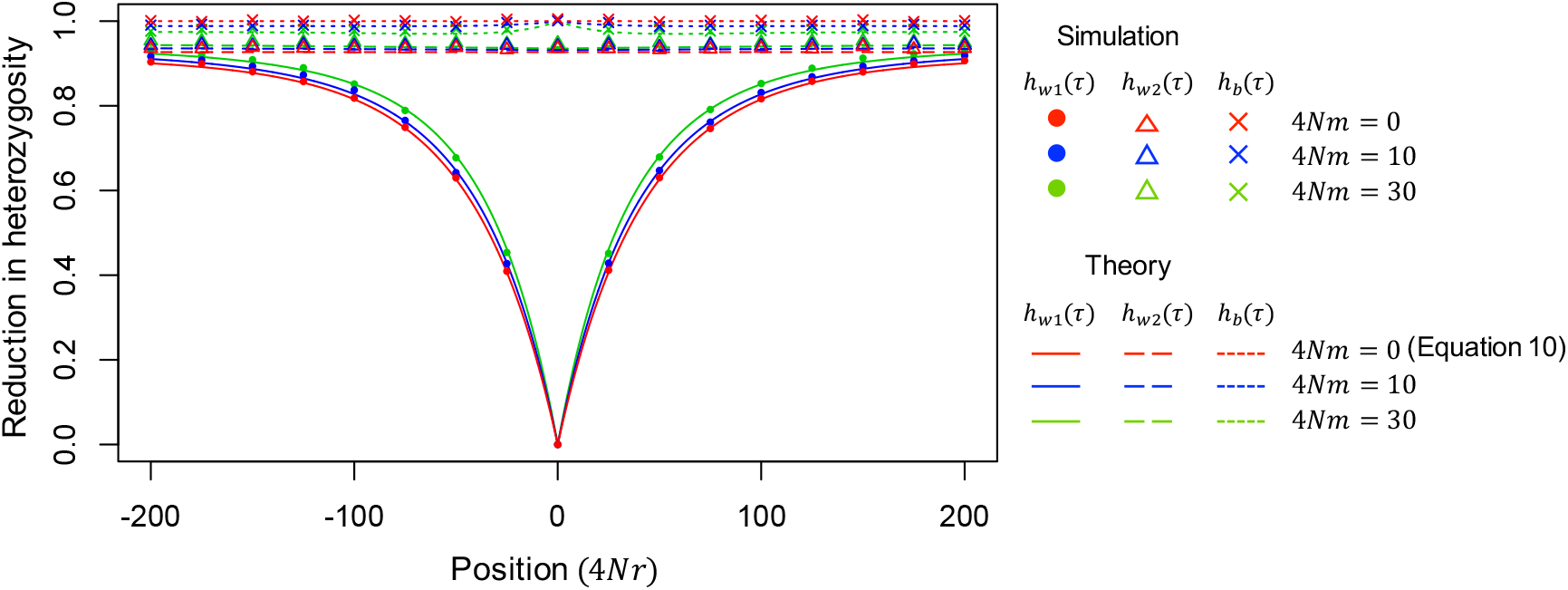
The expected reduction of heterozygosity after a sweep in the two-population model. Position is shown in 4*Nr* from the selected site. *N*_1_ = *N*_2_ = 1000 and *m* = *m*_1_ are assumed. Theoretical results for *h*_*w*1_(*τ*) *h*_*w*2_(*τ*) and *h*_*b*_(*τ*) computed from 13-16 by assuming *y*_*a*1_(0) = *y*_*a*2_(0) = 0.3 for convenience, but very similar results were obtained for other values of *y*_*a*1_(0) and *y*_*a*2_(0). In the case of no migration (red), our results is identical to Stephan *et al.* (1992) (i.e., Equation 10)

### Erosion and growth of a genomic island

When a new locally adaptive mutation (a→A) quasi-fixes in subpopulation I, a block of region in which genetic variation in subpopulation I is dramatically reduced arises (Figure 1B), which is referred to as an initial genomic island. In this section, by using the two-locus model defined in Figure 2, we consider the process after this state, but our derivation is flexible enough to plug in any initial state.

We use a similar diffusion approach to the previous section but we focus on the behavior of *y*_*A*1_ and *y*_*a*2_. The expected changes of an arbitrary function *f* (*y*_*A*1_, *y*_*a*2_) is described as the following ordinary differential equation:

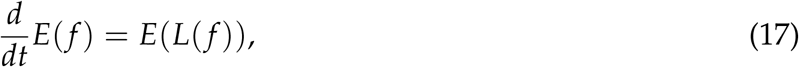

where *L* is a differential operator of the Kolmogorov backward equation, which is given by

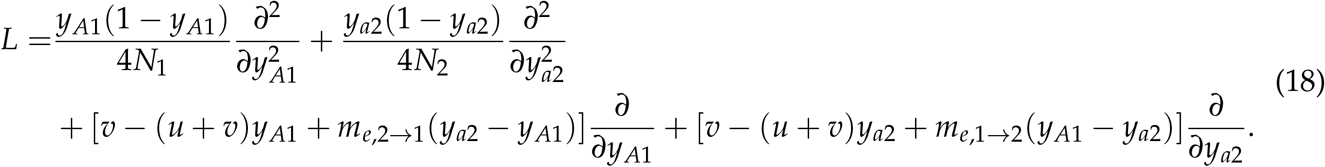

As well as the previous section, we use the effective migration rate (Bengtsson 1985):

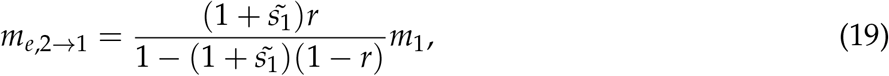

where 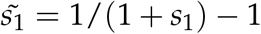 is the relative selection coefficient of maladapted individuals in subpopulation I. *m*_*e*,1→2_ is defined by Equation 14. We consider the dynamics of the first and second order moments, and put 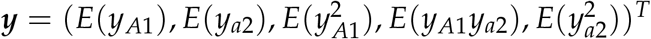. By using Equation 17, we derive a differential equation for ***y*** as follows:

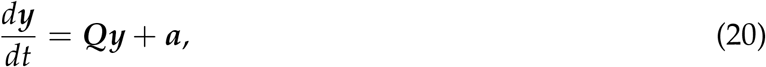

where ***Q*** is the 5 × 5 matrix given by

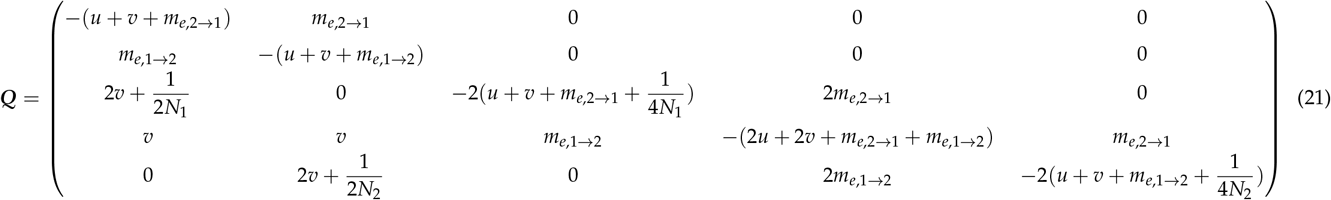

and ***a*** = (*v,v*,0,0,0)^*T*^. By solving Equation 20, ***y*** is given by

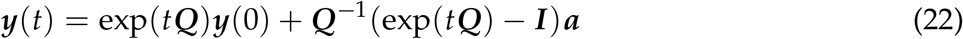

where ***I*** is the identity matrix of size 5. ***y*** at equlibrium is given by 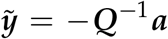. Our solution at equilibrium is well consistent with previous studies (Charlesworth *et al.* 1997; Yeaman *et al.* 2016) that used the coalescent approach (see APPENDIX B).

Figure 5 compares our theoretical results from Equation 22 (broken lines) with simulation results. *N*_1_ = *N*_2_ = 1000, *s*_1_ = −*s*_2_ = 0.05, *u* = *v* = 2.5 × 10^−6^, *m*_1_ = *m*_2_ = 1.25 × 10^−3^ are assumed. As the initial condition (*t* = 0), we set *h*_*w*1_ = 0, *h*_*w*2_ = 0.18 and *h*_*b*_ = 0.1, representing a situation after a local sweep in subpopulation I. Equation 22 describes how a sharp peak of divergence grows along time. As time goes, *h*_*w*1_ and *h*_*w*2_ become closer to each other, and eventually reaches their equilibrium values (t≫10,000). *h*_*b*_ also decreases except for a short region surrounding the selected site. The rate of erosion (decrease of *h*_*b*_) is high as going apart from the selected site. At the selected site, *h*_*b*_ gradually increases and eventually develops a sharp peak. It reaches an equilibrium after a significant amount of time, where the selection-migration balance holds so that the shape of the peak does not change much.

**Figure 5.**
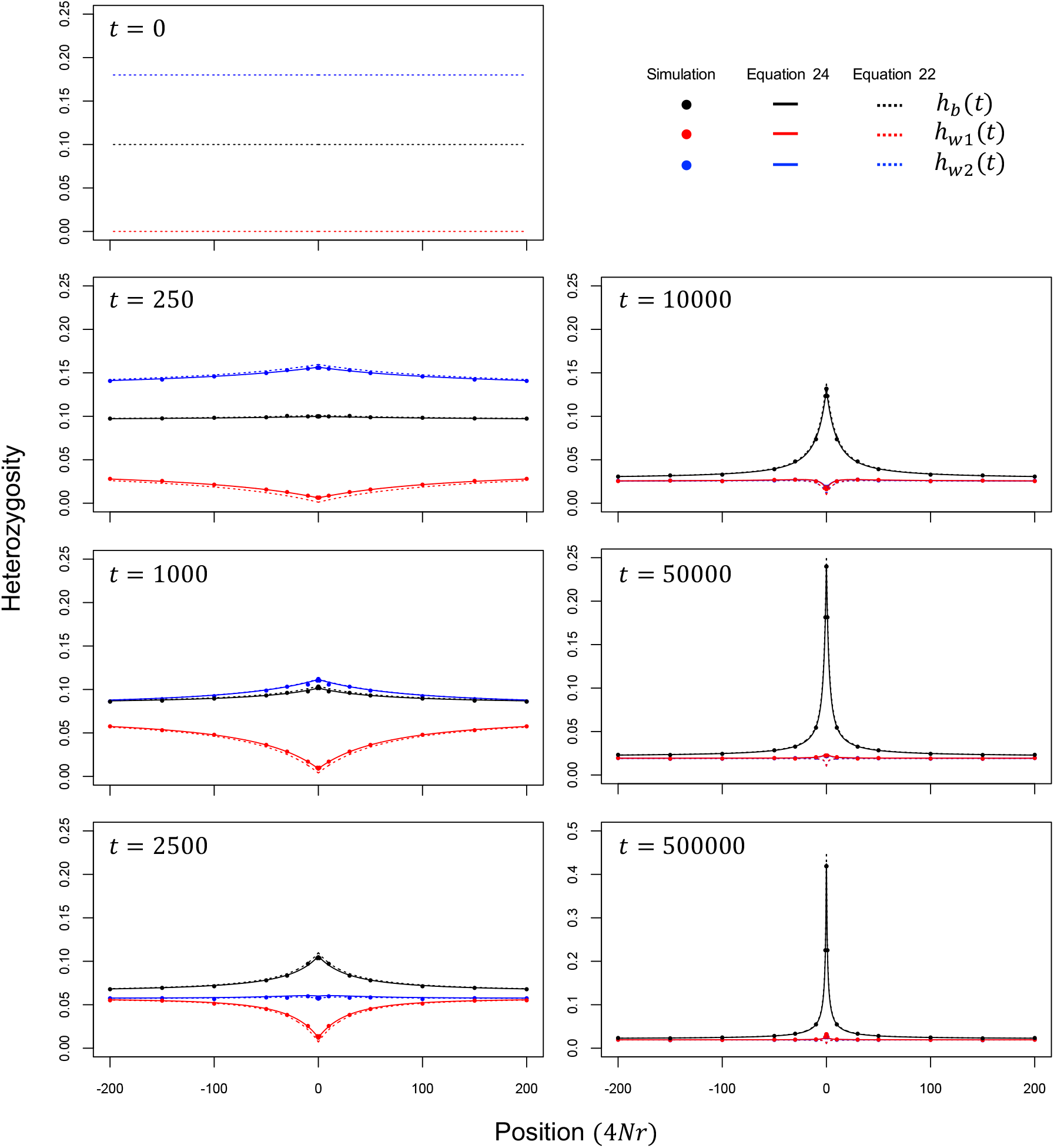
Temporal change of heterozygosity (*h*_*w*1_, *h*_*w*2_, *h*_*b*_) after a local sweep in subpopulation I. Position is shown in 4*Nr* from the selected site. *N*_1_ = *N*_2_ = 1000, *s*_1_ = −*s*_2_ = 0.05, *u* = *v* = 2.5 × 10^−6^, *m*_1_ = *m*_2_ = 1.25 × 10^−3^, *y*_1_(0) = 0.0 and *y*_2_(0) = 0.1 are assumed. Theoretical results from Equations 22 and 24 are shown by broken and solid lines, respectively. Simulation results (closed circles) are the averages over 50,000 replications of forward simulation.

Figure 5 shows that Equation 22 is well consistent with the simulation results (broken lines), but they could be further improved if we include the effect of maladapted alleles, which were completely ignored in Equation 22. In practice, although their effect on the long-term dynamics is ignorable, they stay in the population, thereby constituting a certain proportion; their expected frequencies in subpopulations I and II are, respectively, 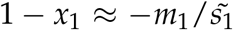 and *x*_2_ ≈ −*m*_2_/*s*_2_. Let us focus on the trajectory of the frequency of a single maladaptive migrant. We ask how long such a maladaptive migrant can survive (as a maladaptive allele). It could be eliminated by selection, or recombine with an adaptive allele and escape from the maladaptive state. Because the expected time until a migrant dies or recombines in subpopulations I and II are, respectively, given by

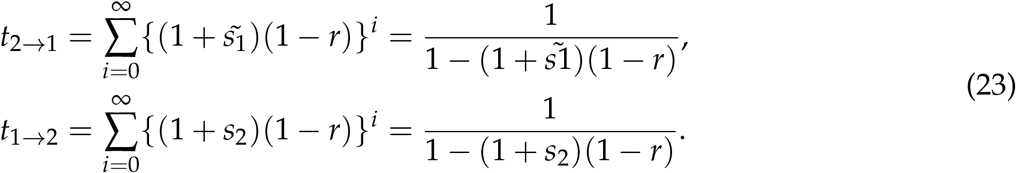

Therefore, the expected numbers of neutral alleles from the other subpopulation with the maladapted allele is *N*_2_*m*_2_*t*_1→2_ and *N*_1_*m*_1_*t*_2→1_ in subpopulations I and II, respectively.

Let the frequencies of B in subpopulations I and II including maladapted ones are denoted by 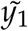 and 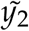. Together with this effect of maladaptive alleles, the first and second-order moments of 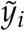 are given as follows,

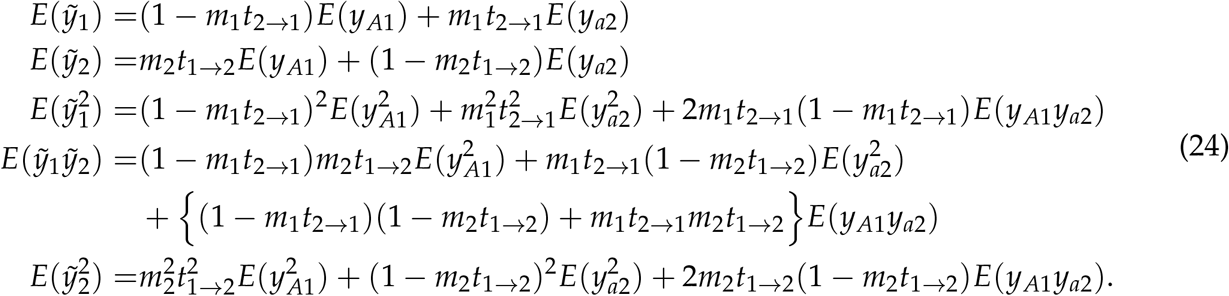

Figure 5 shows that Equation 24 fits to the simulation results better than Equation 22.

## DISCUSSION

In the earlier stages of ecological speciation with gene flow, divergent selection should work to maintain phenotypes that are adaptive to each niche (Wu 2001; Turner *et al.* 2005; Nosil 2012). Therefore, it is predicted that genetic variations responsible to those adaptive phenotypes should appear as genomic islands of speciation. This article theoretically considers the evolutionary behavior of a genomic island of speciation, from its initial establishment to stable preservation. The process was divided into three phases, the establishment, erosion and equilibrium phases (Figure 1). We obtained (i) the establishment probability of a locally adaptive mutation, (ii) the expected reduction of genetic variation within subpopulations I and II after a local sweep that creates an initial genomic island, (iii) the expected erosion of the initial island as a function of time since the sweep, and (iv) the expected shape of the peak of divergence in the island in equilibrium.

For (i), we have successfully derived a close-form formula of the establishment probability along the formulation of Barton (1987). Our simulation showed that our theoretical results for *u*(1/2*N*_1_, 0) and *u*(0, 1/2*N*_2_) outperform the previous studies, although Yeaman and Otto (2011)’s heuristic approach is almost as good as ours. It would be intriguing to discuss the analogy between our result and those of Gavrilets and Gibson (2002) and Whitlock and Gomulkiewicz (2005). Because this work focuses on divergent selection so that allele A is quasi-fixed in subpopulation I whereas allele a is quasi-fixed in subpopulation II, we assume *s*_1_ > 0 and *s*_2_ < 0. However, as showed in Figure 3, it is possible that either A or a could fix in the entire population even if *s*_1_ > 0 and *s*_2_ < 0 hold, although it might take an extremely long time. In contrast, Gavrilets and Gibson (2002) and Whitlock and Gomulkiewicz (2005) obtained the probability of such eventual fixation in the entire population. These studies and ours can be understood in a single framework as follows. Assuming *s*_1_ > 0 and *s*_2_ < 0, the establishment of A first occurs and maintained quite stably for a long time, but with time going infinity, allele A could fix in the entire population most likely when the average selection coefficient 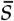 is positive, while allele a could likely fix when 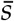 is negative. This is why our formula of the establishment probability (Equation 2) is the same as the numerator of the fixation probability when 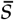 is positive (Equations 7 and 8 in Gavrilets and Gibson 2002 and Equation 6 in Whitlock and Gomulkiewicz 2005). On the other hand, the establishment probability significantly differs from the fixation probability of Gavrilets and Gibson (2002) and Whitlock and Gomulkiewicz (2005) when 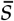 is negative because such a mutation hardly goes to eventual fixation, although it can be maintained as a quasi-fixed state for a sufficiently long time.

For (ii), we extended the diffusion method of Stephan *et al.* (1992) to our two-population model. Because the beneficial allele A fixes only in one subpopulation, the process is very similar to that of a single population model (Stephan *et al.* 1992), except that migration between two subpopulations has some effect. Our theoretical result (see Figure 4) demonstrated a relatively minor effect of migration; with an increasing migration rate, the level of polymorphism in subpopulation I increases because migration brings genetic variation from subpopulation II.

For (iii) and (iv), we considered the erosion of an initial island created by a local sweep, followed by the development of a stable island at equilibrium. This process from erosion to equilibrium can be described by a single formula 22. Furthermore, Equation 22 is flexible enough to plug in any initial state, such as a secondary contact of already diverged subpopulation. To demonstrate this, in Figure A1, we compare the pattern after a local sweep (left panels) and that after a secondary contact (right panels). After a secondary contact, *h*_*b*_ is already high across the genome, and *h*_*b*_ gradually decreases but selection works to keep divergence around the selected site, thereby creating a peak of divergence (i.e., island). After a very long time (i.e., in equilibrium), the shape of the peak becomes identical to that after a sweep.

We have thus developed analytical expressions for the evolutionary behavior of genomic island of speciation, from the emergence of an initial island by a local sweep to stable maintenance of the island in equilibrium. Genomic islands of speciation can arise in the earlier stages of ecological speciation, but it does not necessarily mean that the emergence of genomic islands of speciation always results in speciation. It is possible that genomic islands of speciation could disappear by environmental changes or by chance, and no speciation occurs. To achieve speciation, there would be many other forces necessary, including emergence of additional islands (Feder *et al.* 2012a; Via 2012; Feder *et al.* 2012b; Aeschbacher and Bürger 2014; Yeaman *et al.* 2016), further divergence on a genomic-scale possible due to a reduction in migration rate, and environmental changes. More theoretical works are needed to fully understand the process to ecological speciation.

## APPENDIX

### Appendix A: The solution of Equations 4 and 5

First, we present a proof that there is at most one solution which fulfills *p*_1_ > 0 and *p*_2_ > 0, and the condition on which such a solution exists is *a* + *d* > 0 or *ad* – *bc* < 0. Then, we give a closed expression of the solution.

For *ψ*_1_ and *ψ*_2_ to satisfy *p*_1_ > 0 and *p*_2_ > 0, *ψ*_1_ > 0 and *ψ*_2_ > 0 are needed. Notice that *b, c* > 0 because migration rate and population size are always positive. We put *f* (*x*) = *x*^3^ – 2*ax*^2^ + (*a*^2^ – *bd*)*x* + (*abd* – *b*^2^*c*) and *f* ^*t*^(*x*) = 3*x*^2^ – 4*ax* + (*a*^2^ - *bd*).

1. *a* ≥ 0 From Equation 5, *ψ*_1_ > *a* is needed. Because *x*-cordinate of vertex of 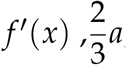, is not greater than *a, f*′ (*x*) monotonically increases when *x* > *a*. Noting *f* (*a*) = −*b*^2^*c* < 0, there is only one solution.
2. *a* < 0 and *d ≤* 0 From Equation 5, *ψ*_1_ > 0 is needed. Because *f*′ (0) = *a*^2^ - *bd* > 0 and the *x*-coodinate of vertex of 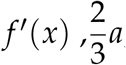, is smaller than 0, *f*′ (*x*) > 0 when *x* > 0. Therefore, whether *f* (*x*) = 0 has a solution or not in (0, ∞) depends on the sign of *f* (0). If *f* (0) ≥ 0, i.e. *b*(*ad* – *bc*) ≥ 0, there is no solution. Otherwise, there is only one solution.
3. *a* < 0 and *d* > 0 From Equation 5, *ψ*_1_ > 0 is needed. Because *x*-coodinate of vertex of 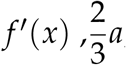, is smaller than 0, *f*′ (*x*) monotonically increases when *x* > 0. Noting *f* (0) = *abd* – *b*^2^*c* < 0, there is only one solution.

Noting that *ad* – *bc* is negative when *ad ≤* 0, the condition on which one solution exists is reduced to *a* + *d* > 0 or *ad* – *bc* < 0. This is the same as the condition where a deterministic model,

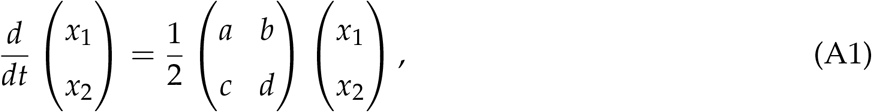

has a positive growth rate, in other words, the matrix in Equation A1 has at least one positive eigenvalue.

Next, we present a closed form of *ψ*_1_. From the above proof, if there is a nonzero real root of *f* (*ψ*_1_) = 0 which fulfills *p*_1_ > 0 and *p*_2_ > 0, the root is the largest real root of *f* (*ψ*_1_) = 0. Therefore, by using the solution of cubic equation, *ψ*_1_ can be expressed as

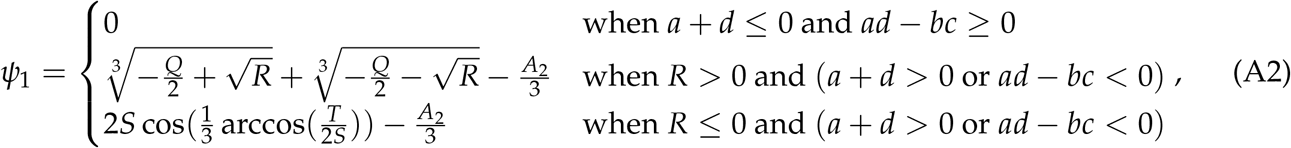

where 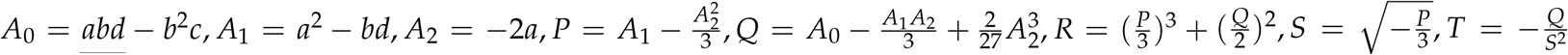. In the above expression, we assume the range of principal value of *y* = arccos(*x*) as 0 *≤ y ≤ π*.

### Appendix B: Comparison between diffusion and coalescent at equilibrium phase

In the main text, we show that replacing the migration rate in the neutral diffusion equation by the effective migration rate well approximates the effect of linkage with the locus under divergent selection. In a neutral model, heterozygosity at equilibrium in a structured population is already well studied by the coalescent theory under the infinite-site model (reviewed in Wakeley 2009). In this work, we alternatively used the forward diffusion approach because the diffusion approach can be applied to more general conditions. In this Appendix, we show our diffusion result at equilibrium is consistent with that of the coalescent theory.

We attempt to derive the expected heterozygosity under the infinite-site setting along our diffusion-based derivation. In practice, we first consider a *K*-allele model, and then the results will be transformed to the infinite-site model. Let B allele be one of the alleles at the locus. We put *y*_1_ and *y*_2_ as frequency of allele B in subpopulation I and II, respectively. In the following derivation, we assume *N*_1_ = *N*_2_ = *N* and *m*_1_ = *m*_2_ = *m*. The differential operator of the Kolmogorov backward equation is as follows,

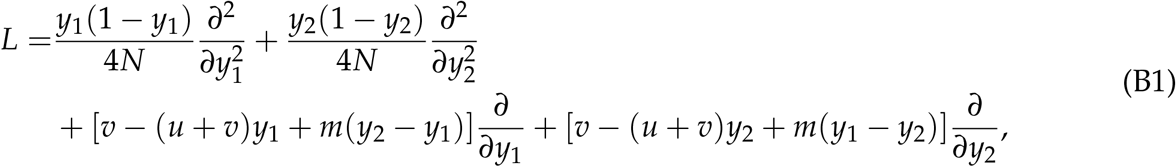

At the equilibrium, we derive the moments up to the second order as

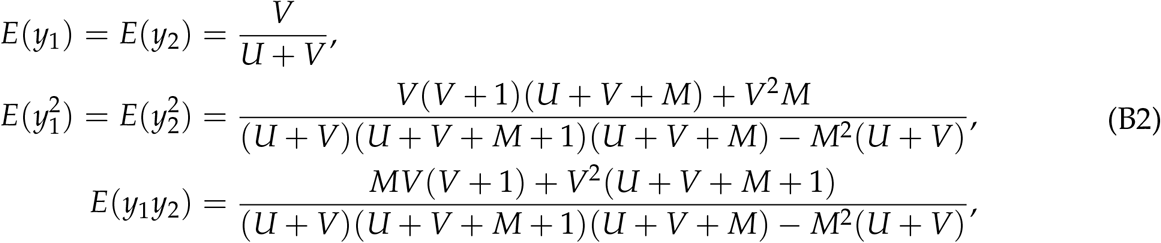

where *U* = 4*Nu, V* = 4*Nv* and *M* = 4*Nm*. In the limit to the infinite-allele model, that is, 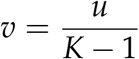 and *K* → ∞, the expected heterozygousity within and between subpopulation goes to

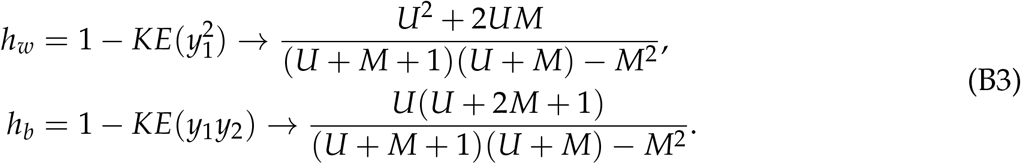

This result udner the infinite allele-setting can be transformed to the infinite-site mode: If we put 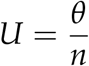 and *n* goes to ∞, *π*_*w*_ and *π*_*b*_ are described as

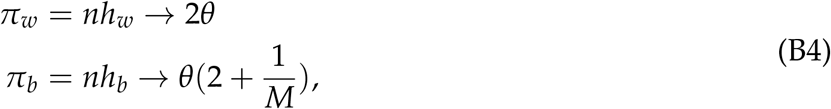

which is identical with the result from the coalescent theory (Charlesworth *et al.* 1997; Yeaman *et al.* 2016).

### Appendix C: Equation 22 for a scenario of secondary contact

We compute Equation 22 for a scenario of secondary contact, where we assume that already diverged two subpopulation have merged so that there are a number of fixed sites between the two subpopulations. To make a realization of this situation, we set *y*_1_(0) = 0.1 and *y*_2_(0) = 0.9, and the other parameters are identical to those used for Figure 5. Figure A1 compares the patterns after a local sweep (left panels) and after a secondary contact (right panels). After a secondary contact, *h*_*b*_ is already high across the genome, and *h*_*b*_ gradually decreases but selection works to keep divergence around the selected site, thereby creating a peak of divergence. In equilibrium, the shape of the peak becomes identical to that after a sweep.

**Figure A1.**
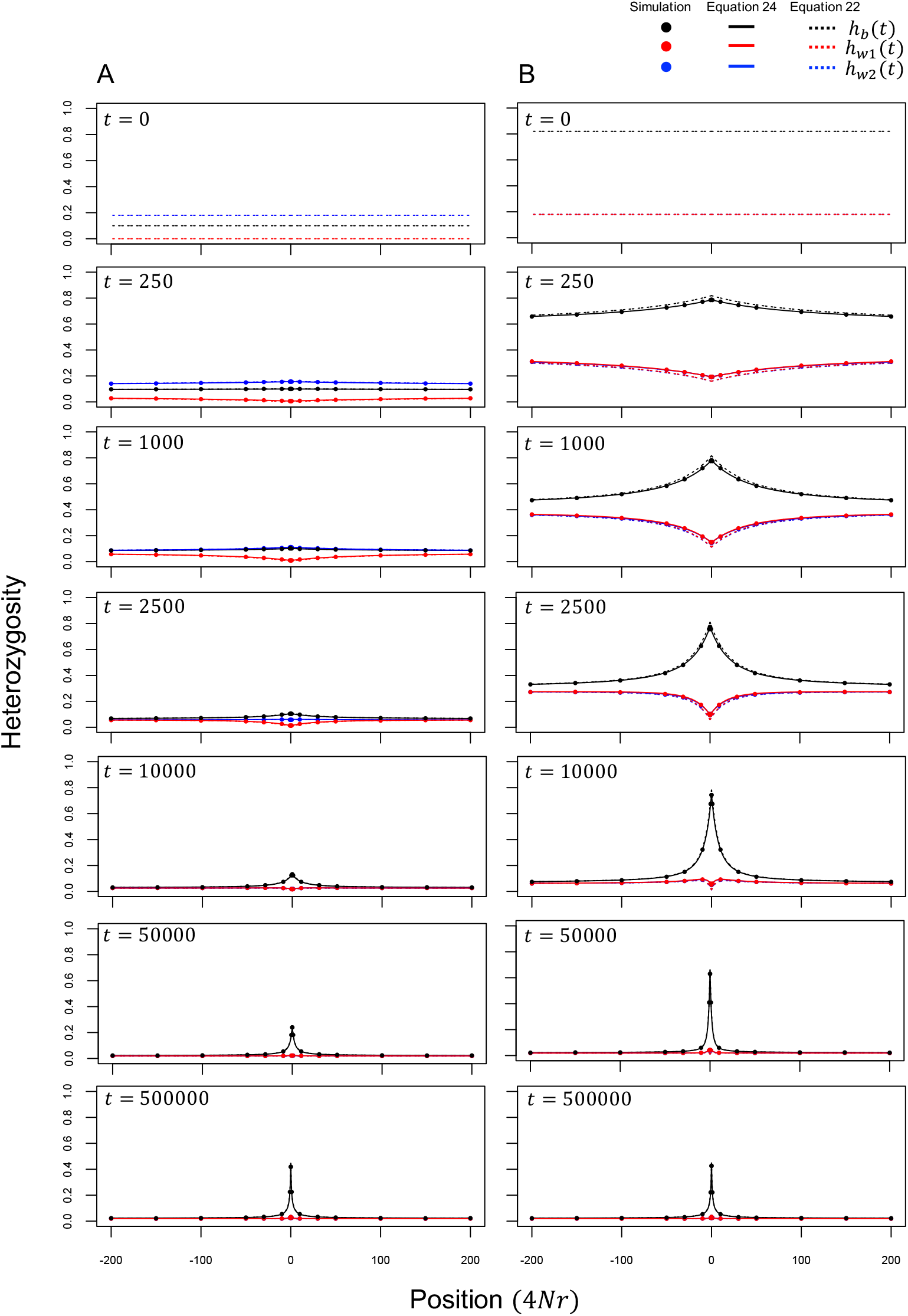
Temporal change of heterozygosity (*h*_*w*1_, *h*_*w*2_, *h*_*b*_) as a function of recombination rate (A) after a local sweep in subpopulation I and (B) after a secondary contact. *y*_1_(0) = 0.0 and *y*_2_(0) = 0.1 are assumed in (A), whereas *y*_1_(0) = 0.1 and *y*_2_(0) = 0.9 in (B). Theoretical results from Equations 22 and 24 are shown by broken and solid lines, respectively. Simulation results (closed circles) are the averages over 50,000 replications of forward simulation. The left panel is identical to Figure 5.

